# *Drosophila* males require the longitudinal stretch receptors to tremulate their abdomen and produce substrate-borne signals during courtship

**DOI:** 10.1101/2024.05.13.593852

**Authors:** Jonathan K. M. Lee, Eugenie C. Yen, Caroline C. G. Fabre

## Abstract

Substrate-borne cues are important species-specific signals that are widely used during courtship of many animals, from arthropods to vertebrates. They allow mating partners to communicate with, recognise and choose one another. Animals often produce substrate-borne signals by vibrating a body part, such as the abdomen. During *Drosophila* courtship, species-specific substrate-borne vibrations are generated by the male’s regular up-and-down abdominal tremulations and these must be precisely controlled to produce an effective and specific signal. The vibrations immobilise the female, therefore facilitating copulation. It is not known how the male’s nervous system regulates this abdominal tremulation. Here, we demonstrate a role for the dorsal abdominal longitudinal stretch receptors (LSR), which include the dorsal bipolar dendritic (dbd) neurons. These neurons are a set of conserved proprioceptors found throughout Insecta. We show that impairing the function of dbd neurons through general inhibition results in males exhibiting high level of arhythmic abdominal movements (referred to as bobbing) and decreased level of tremulation. Strikingly, this causes a failure in the females’ response during courtship. We show that depleting the mechanosensitive ion channel TRPA1 (but not Piezo) in the dbd neurons leads to a similar increase in bobbing movements. Thus, we identify neurons and a key molecular player necessary for males to perform this important mode of communication.

## Introduction

In addition to visual, acoustic and chemical cues, many arthropod and vertebrate groups use substrate-borne vibratory signals to communicate with conspecific individuals, such as during mating, predation and parental care^1–3^. Abdominal vibratory movements (also called abdominal tremulations) are one way animals generate substrate-borne signals^1,3^. For example, the males of Salticid spiders, plecopteran, pentatomid bugs, and red mason bees, all produce substrate-borne vibrational signals by tremulating their abdomen during courtship, influencing the receptivity of the female^4–6^. Tremulations produce species-specific vibrations in the substrate^1,3^. As such, the regulation of such stereotyped vibratory movements must require interplay between the central nervous system (CNS) and the peripheral nervous system (PNS). It remains unknown, however, which central and peripheral neuronal circuits, and the genes within, control these abdominal movements.

We showed that *Drosophila melanogaster* males use abdominal tremulations during courtship^7^. These tremulations occur during ∼30% of the courtship displays and consist of up-and-down oscillations in the form of bouts lasting between 1s and 3s with a pulse repetition rate of around 5Hz^7^. These signals immobilise the female, a response which facilitates copulation^7–9^. Sensory feedback is important to shape rhythmic behaviours^10^. In *Drosophila* the larval body wall multidendritic (md) sensory neurons act in the regulation of rhythmic locomotor patterns in larvae^11–15^. Some of these neurons persist throughout metamorphosis and are present in the adult abdomen, albeit with remodelled neural processes^16–18^. This remodelling likely reflects their roles at each developmental stage; however, few reports provide information about their function in adults^19–21^. It is therefore possible that these neurons have a role in adult rhythmic behaviours, such as the tremulations. Here, we present our results for a subset of md neurons, the persistent dbd (dorsal bipolar dendritic) neurons localised in the dorsal hemisegments (the tergite) of the abdomen. dbd neurons have a unique sensory innervation pattern and develop in close contact with a dorsal longitudinal muscle in each abdominal hemisegment, forming the conserved Longitudinal Stretch Receptor (LSR)^22–24^. First, we manipulated the function of these neurons using a specific Gal4 driver line combined with the effector construct UAS-shibire^ts^ that transiently inhibits the traffic of neurotransmitters in the neurons targeted at restrictive temperature^25,26^. We found that this led to abnormalities in the display of abdominal movements during male courtship, with subsequent disruptions in the female’s response, an effect that was not observed when disturbing a different set of tergite md neurons, the (larval class IV) ddaC neurons. We characterised the fly adult LSR organs further by identifying aspects of their expression patterns and anatomy, and by analysing a driver line that specifically targets them. By expressing RNAi constructs that knock-down the expression of the mechanosensory ion channels TRPA1 and Piezo, we identified a role for TRPA1, but not Piezo, in male abdominal movements during courtship. Our results suggest that dbd neurons, via TRPA1 ion channels, act as muscular stretch sensors that sense the abdominal oscillations and send this information on to central targets for abdominal movement regulation during courtship.

## Results

### Perturbing Clh8-Gal4-expressing neurons increases bobbing and reduces tremulations in courting males, leading to a reduced female response

We previously reported that *Drosophila melanogaster* males perform bouts of regular up-and-down abdominal tremulations during courtship, which produce substrate-borne vibrations that are simultaneous with female pausing^7^. These abdominal tremulations have been described in the wild type and under several genetic and environmental variables^7,27^. Looking anew at *Drosophila melanogaster* wild types males, we noticed another, rarer, type of abdominal movement called bobbing that was previously reported only in *Drosophila simulans* and *Drosophila erecta* males^28^. Bobbing consists of jerky up-and-down abdominal movements of large amplitude, either singly or several consecutive bobs^28^. *Drosophila melanogaster* Oregon R (OrR) wildtype flies displayed a rate of ∼3 ± 0.5 bobs per minute (Figure 1A). These bobbing behaviours did not coincide with any specific behaviour or response in the courted female, and only ∼9 ± 4 % of the bobbing movements coincided with the paired female being immobile (135 bobs analysed over 10 pairs of OrR flies). Besides, these bobbing movements did not produce any detectable acoustic or substrate-borne vibrations, as observed with our microphone or laser vibrometer, respectively, suggesting they may not offer a form of communication during courtship.

**Figure 1.**
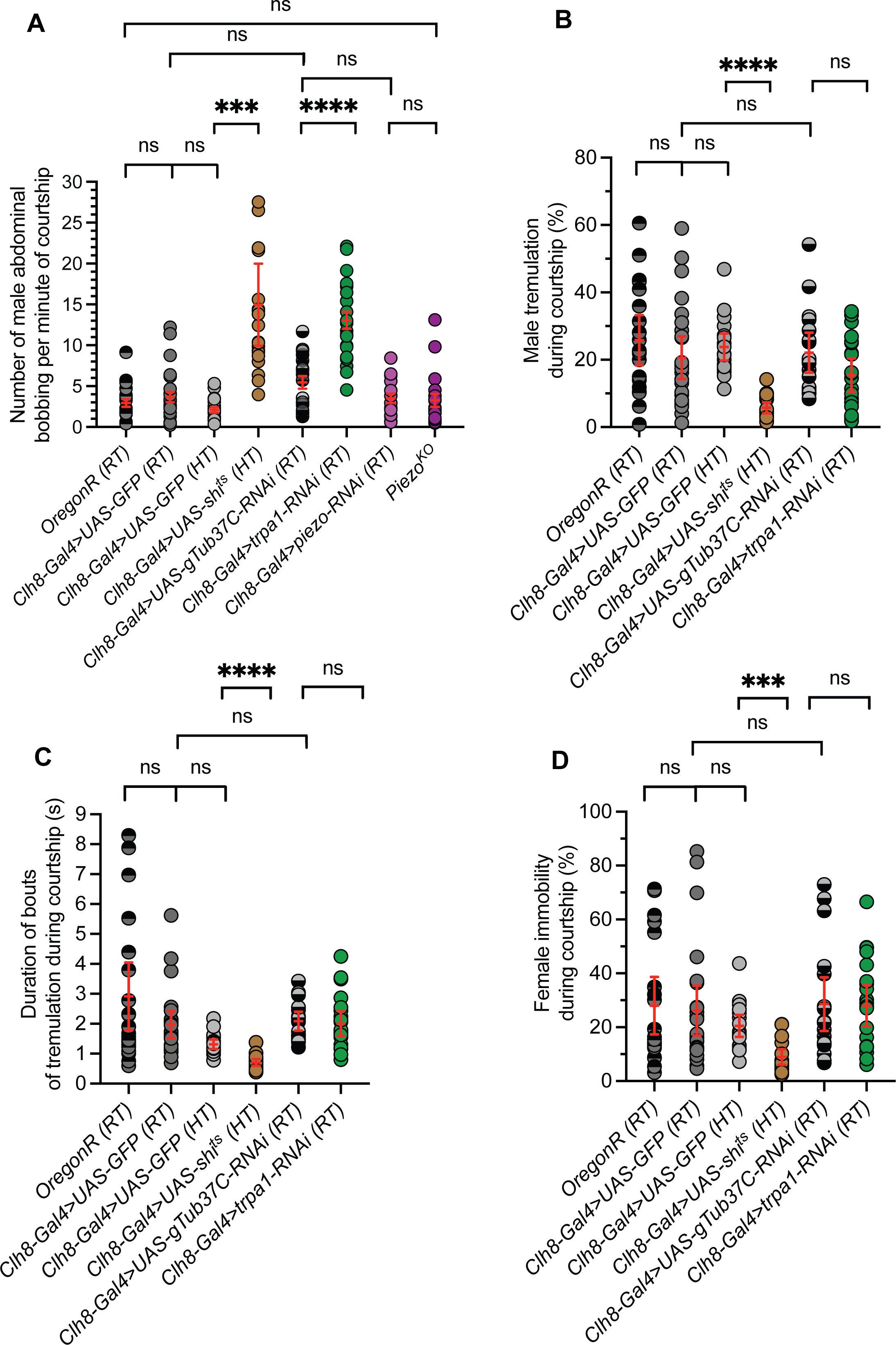
Quantification of experimental and control male and female behaviours during courtship. In all pairs females are wild-type OregonR. Pairs were monitored at room temperature (RT) or high temperature (HT) depending on experimental requirement. **(A)** Number of abdominal bobbing displayed by courting males is shown per minute of courtship. Data is shown for control pairs including OregonR males (column1 at RT, black/grey) or *Clh8-Gal4>UAS-GFP* males (column 2 at RT, dark grey; column 3 at HT, light grey), to compare with pairs including the experimental males carrying *Clh8-Gal4>UAS-shi^ts^* (column 4 at HT, brown). Data is also shown for control pairs including *Clh8-Gal4>UAS-gTub37C-RNAi* males (column 5, black/light grey) to compare with pairs including experimental males carrying *Clh8-Gal4>UAS-trpa1-RNAi* (column 6, green) or *Clh8-Gal4>UAS-Piezo-RNAi* (column 7, magenta). Finally, data is shown for pairs including a *piezo^KO^* male (column 8, purple) to compare with OregonR controls. Ethograms were constructed from analysis of 19, 21, 19, 21, 19, 21, 16, and 18 pairs (in the order illustrated on the graphs). The number of male bobbing increased significantly in pairs monitored at HT that included a male *Clh8-Gal4>UAs-shi^ts^*, and in pairs monitored at RT that included a male *Clh8-Gal4>UAS-trpa1-RNAi*. *Clh8-Gal4>UAS-Piezo-RNAi* and *dpiezo^KO^* males behaved similarly to control. **(B)** The percentage of time males display tremulation during courtship is shown. Same pairs as in (A) with similar colour coding of genotypes. The percentage of time *Clh8-Gal4>UAS-shi^ts^* males displayed tremulation during courtship is significantly reduced compare with controls. **(C)** The average duration of individual bouts of tremulation is shown for the same pairs as in (B). Bouts of tremulations performed by courting *Clh8-Gal4>UAS-shi^ts^* males (HT) are significantly shorter than those performed by control males. **(D)** The percentage of female immobility during courtship is shown for the same pairs as (B). Females remain significantly less immobile when courted by *Clh8-Gal4>UAs-shi^ts^*male.

To identify neurons involved in tremulations, we screened through various Gal4 lines expressing the thermosensitive *shibire* effector (*UAS-shi^ts^*)^25^ within different subsets of abdominal sensory neurons in courting males paired with OrR females. Strikingly, at the restrictive temperature (30°C), expressing *UAS-shi^ts^* with the line *Clh8-Gal4*^11^ led to a large increase in the occurrence of bobbing, where males displayed ∼15 ± 2 bobbing movements per minute during courtship (Movie S1), while controls both at permissive (23°C) and restrictive temperatures displayed only ∼4 ± 0.6 and ∼2 ± 0.3 bobbing movements per minute, respectively (Figure 1A; *p=0.0041* and *p=0.0008*, respectively). When placed at the restrictive temperature on their own (i.e. not mixed with a female), *Clh8-Gal4>UAS-shi^ts^* males did not perform bobbing (Movie S2), suggesting that increased bobbing occurs only during courtship. Control males displayed tremulations for ∼21 ± 3 % and ∼24 ± 2 % of the courtship time (at permissive and restrictive temperatures, respectively) (Figure 1B). We previously showed that bouts of tremulations in the wild type OrR showed an average duration of ∼3 ± 0.5 seconds^7^, a result which we recapitulated with similar values for our experimental controls (Figure 1C); we found that “short” bouts of tremulation with a duration lower than 1s only represented ∼27 ± 1 % of the total number of tremulation bouts in controls (N=255 bouts, 11 flies). However, *Clh8-Gal4>UAS-shi^ts^* males showed a large reduction in the occurrence of tremulations, where males displayed only ∼6.5 ± 1 % of tremulation per courtship display (*p=0.0010* and *p<0.0001* when compared with controls; Figure 1B), during which short bouts represented ∼80 ± 3 % of the tremulation bouts (*p=0.002*; N=185 bouts, 7 flies); the average duration of individual bouts of tremulation displayed by courting *Clh8-Gal4>UAS-shi^ts^* males was significantly shorter than controls’, with a value below 1s at ∼0.7 ± 0.05 s (Figure 1C).

We and others also previously showed that tremulations generate substrate-borne vibrations with characteristic interpulse intervals (IPIs) at specific temperatures^7,27,29^. Laser vibrometry recordings of courting and tremulating *Clh8-Gal4>UAS-shi^ts^* males showed that the IPIs of their substrate-borne signals exhibited similar IPI values to controls at the restrictive temperature (Supplementary Figure S1A). Thus, these flies still produce normal signals during tremulation, but the length and occurrence of the tremulation bouts are reduced. Besides, we did not observe defects in any of the other courtship behaviours of *Clh8-Gal4>UAS-shi^ts^* males with females. For example, their wing fluttering that produce the acoustic serenade^30^ were displayed at similar levels to control (Supplementary Figure S1B).

To test whether the increased occurrence of bobbing and reduced occurrence and length of tremulation bouts affected female receptivity, we monitored the immobility of wild-type OrR females paired with *Clh8-Gal4>UAS-shi^ts^* males at restrictive temperature. We found that female immobility was globally reduced compared to controls (*p=0.0004* at restrictive temperature for instance, Figure 1D), and we further found that the specific immobility response to tremulations was significantly reduced (*p<0.0001*; Supplementary Figure S1C). We tested whether this reduction in the female response was directly related to shortened tremulation bouts by comparing the response of females to “short bouts” (<1s) versus long bouts (>1s). In both control pairs and pairs with *Clh8-Gal4>UAS-shi^ts^* males, only long tremulation bouts were associated with female immobility (Supplementary Figure S1D). Thus, the duration of individual tremulation bouts appears to be an important factor in triggering female immobility during courtship.

### Dbd neurons are the Clh8-Gal4-expressing neurons necessary for proper regulation of male abdominal movements during courtship

We next wanted to identify which adult male neurons must express *Clh8-Gal4* in order to regulate bobbing and tremulations. In larvae, *Clh8-Gal4* is expressed in the longitudinal (dbd, vbd) and lateral (lbd) bipolar dendrite neurons, and in the dmd1 neurons^11^. Only dbd neurons survive in adults^18^ and we observed *Clh8-Gal4>UAS-mGFP* expression in these neurons, which localise to the dorsal body wall of the abdomen (Figure 2). Development of the dbd neurons during pupal stages and into the pharate adult has been described in detail^18,24^ and we found the anatomy to be similar, even in 3-day-old adults (Figure 2A). In each dorsal hemisegment in adults, the dendrite of the bipolar dbd neuron extends along the dorsal longitudinal muscle that is fifth from the dorsal midline (Figure 2A-B); these muscles are believed to be part of the retractor muscles of the dorsal abdomen^24,31^ and could conceivably be required for tremulations. This combination of dbd neurons with an associated muscle makes up the longitudinal stretch receptors (LSR), with one LSR per adult dorsal abdominal hemisegment^23,32^. LSR are conserved stretch receptors that are found in the abdomen of all investigated insect orders, but whose adult function is unknown. The dbd neuron dendrites span a muscle fiber of the LSR precisely suggesting a role in sensing stretching of that muscle^23,24,32^ (Figure 2).

**Figure 2.**
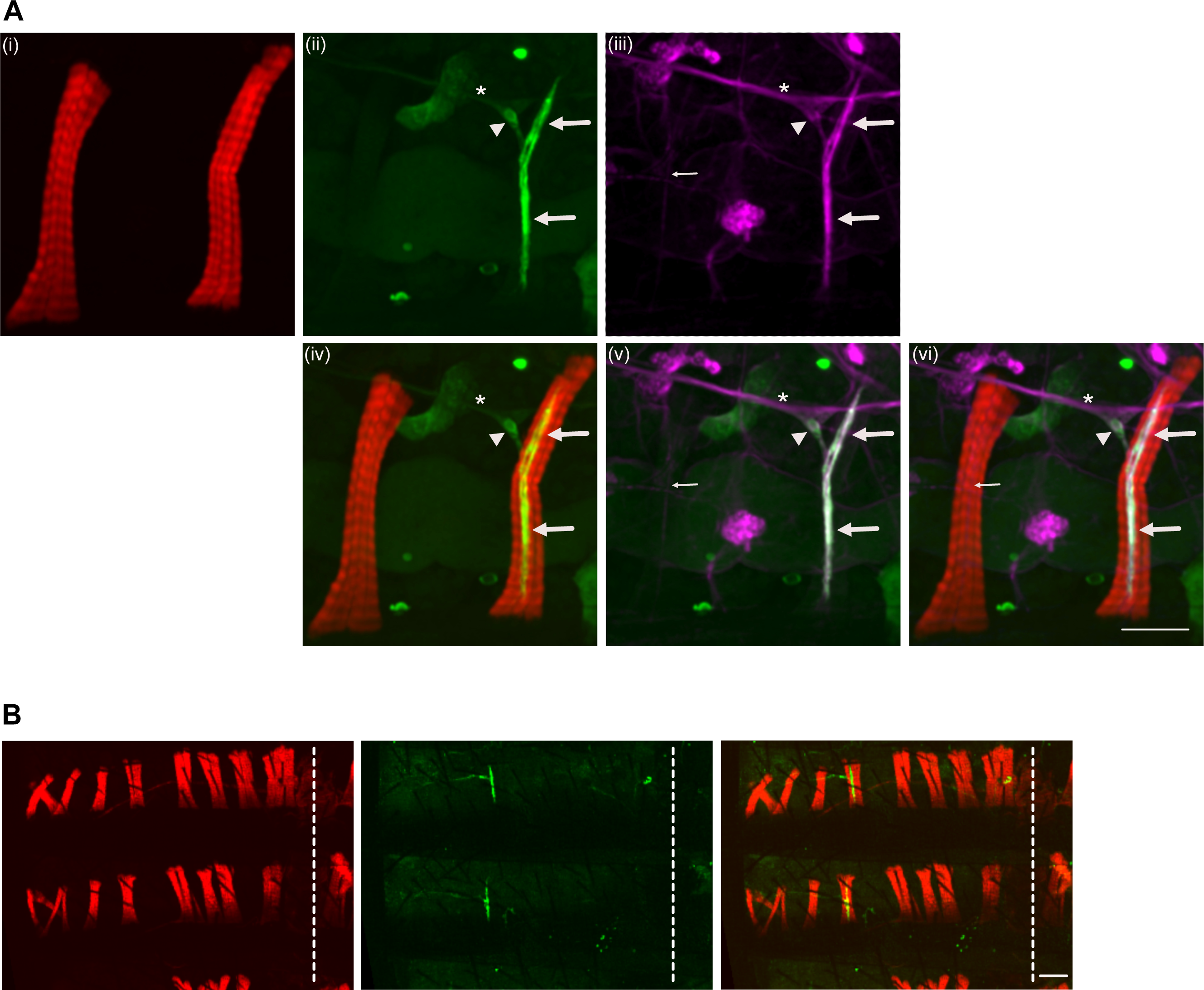
*Clh8-Gal4>UASmGFP* expressing dorsal bipolar dendritic (dbd) neurons are part of the Longitudinal Stretch Receptor (LSR) of adult *Drosophila* abdomen. **(A) Top:** (i) Two longitudinal muscles of the second left hemisegment of the abdomen of a male carrying the constructs *Clh8-Gal4>UASmGFP* are marked with Phalloidin (Red). (ii) Anti-GFP staining in the same tissue reveals the cell body of the dbd neuron (arrowhead), and its bipolar dendrites (large arrows) that are targeted by *Clh8-Gal4>UASmGFP* (green). Axon of the dbd joins the abdominal peripheral nerve of hemisegment 2 (asterix); (iii) anti-HRP staining labels all neuronal membranes (magenta), including the abdominal peripheral nerve on which the dbd axon bundles (asterix), the dbd cell body (arrowhead) and the dbd bipolar dendrites (large arrows); the axons of the motor neurons (small arrow) that innervate longitudinal muscles are visible at the neuromuscular junction of the (left) muscle that is not associated with a dbd neuron. **Bottom:** Merges; (iv) shows merge of (i) and (ii) to show the dbd neuron (green) cell body (arrowhead) and bipolar dendrite (large arrows) extending along the muscle fibers (red) of one of the longitudinal muscles; (v) shows merge of (ii) and (iii) to show neuronal processes that include the dbd neurites; the dbd axon bundles with the nerve (asterix). (vi) Merge of all stainings. LSR neuroanatomy shows it includes the dbd neuron and its associated muscle. The muscle localised to the left does not display such innervation by a sensory neuron. Note, in (ii) and associated merges, green blobs are pieces of auto fluorescent cuticles; in (iii) and associated merges, magenta blobs are unspecific staining of anti-HRP. **(B)** Second and third left hemisegments of a *Clh8-Gal4>UASmGFP* male abdomen are shown, with muscles stained using phalloidin (red), and *Clh8-Gal4>UASmGFP*-expressing neurons stained using anti-GFP (green). One dbd neuron is observed in each hemisegment, each associated with a longitudinal muscle localised 5^th^ from the dorsal midline (dashed line), thus forming together one LSR per hemisegment. Anterior up, Posterior down.

We identified another line, *8-113-Gal4*^11^, which also targets the dbd neurons (Supplementary Figure S2A). Using this line, we could recapitulate the abnormal levels of male bobbing by driving *UAS-shi^ts^* at restrictive temperature (Supplementary Figure S2C). However, using *7-33-Gal4*^11^ to drive expression of *UAS-shi^ts^* in nociceptive sensory ddaC body wall neurons, which localise near the dbd neurons but are not associated with abdominal muscles^18^ (Supplementary Figure S2B), did not lead to an increased occurrence of bobbing (Supplementary Figure S2C). As with larval dbd neurons, we found that the acetylcholine marker (*Cha*-Gal4) drove expression in adult dbd neurons, indicating that dbd neurons express choline acetyltransferase (ChaT) and are cholinergic^33,34^(Supplementary Figure S3A). Consistent with this, using a promoter to express the Gal4 inhibitor Gal80 specifically in cholinergic neurons (*ChaT-Gal80*)^25^ reduced bobbing in *Clh8-Gal4>UAS-shi^ts^* males, while expressing Gal80 specifically in glutamatergic neurons (*VGlut-Gal80*)^35^ had no effect (Supplementary Figure S2B). In summary, the abdominal phenotypes observed in courting *Clh8-Gal4>UAS-shi^ts^* males are due to disturbing the function of the dbd neurons, most likely by impeding their role in stretch-sensing of the abdominal dorsal musculature during courtship.

To determine which gene may be regulating *Clh8-Gal4* expression, and thus may be important for dbd neuron development or function, we aimed to determine the location of *Clh8-Gal4* within the genome. *Clh8-Gal4* was generated during a Gal4 enhancer trap screen by mobilizing an X-linked *P{GawB}* onto chromosomes III^11^. To determine the genomic insertion site of *P{GawB}clh8*, we therefore performed nucleotide BLAST (NCBI) searches using sequences obtained from a Splinkerette PCR (spPCR) assay, which is an adapted version of inverse PCR optimised to identify flanking sequences of P-elements^36^. Using this approach, we identified the neurodevelopmental *furry* gene on chromosome 3L, which showed 99% local sequence similarity (Supplementary Figure S4). Furry is a multi-isoform gene that has previously been implicated in dendritic tiling^37^ and so may be important for the unique dendrite morphology of the dbd neurons along their associated muscle fiber. Further experiments will be required to determine whether adult dbd neurons require the neurodevelopmental gene Furry and precisely what Furry may be doing in these neurons.

### TRPA1, but not Piezo, is required for dbd neurons to regulate tremulation occurrence and bout duration

Stretch receptor neurons require mechanosensory ion channels to mediate their response^38^. We therefore tested whether specific ion channels are required in dbd neurons to regulate abdominal movements during courtship. We considered DmTRPA1 and DmPiezo, two conserved mechanosensory ion channels that are expressed in larval dbd neurons^39^. We found that both were also expressed in adult dbd neurons (Supplementary Figure S5A-B). To test for a role in regulating abdominal movements during courtship, we used *Clh8-Gal4* to drive the expression of RNAi constructs against either DmTRPA1 or DmPiezo in dbd neurons and monitored the behaviour of courting males (Figure 1). As a control, we expressed *UAS-*γ*-tubulin-37C-RNAi*, which targets the maternally-expressed γ*-tubulin-37C* gene, as has been used previously as a negative control for neuronal analyses^40–42^ (Figure 1). Expressing *UAS-DmPiezo-RNAi* did not lead to any increase in bobbing compared to controls (Figure 1A). This result was confirmed by analysing *Dmpiezo* homozygous mutant males that are viable and coordinated^20^ (Figure 1A). In contrast, however, expressing UAS-Dm*TRPA1-RNAi* led to an increase in the occurrence of bobbing events, similar to when expressing *UAS-Shi^ts^* (Figure 1A). Nevertheless, there was no associated reduction in the occurrence or the duration of abdominal tremulations, or the immobility of the female (Figure 1B-D). Knocking down DmTRPA1 therefore leads to an intermediate effect on courting male abdominal movements.

## Discussion

### Sensory feedback provided by the dbd neurons for the regulation of tremulation

Proprioception, the internal sense of body position, affects rhythmic movements, with several examples in vertebrates and invertebrates^43^. We asked which abdominal body wall sensory neurons affect abdominal movements during courtship in the male. We found that manipulating the function of the dbd neurons of LSR organs caused significant defects in the abdominal movements of courting males. Males lacking functional dbd neurons produced higher levels of bobbing, (jerky arrhythmic movements of the abdomen). Some tremulations were displayed in these manipulated males, but of a significantly low occurrence and duration. These findings indicate that sensory feedback from the LSR organs, particularly from the dbd neurons within them, is essential for central circuits to enable limited levels of abdominal bobbing and sustained, consistent tremulations during courtship.

### Circuitry underlying the abdominal movements during courtship

The muscles of the LSR are believed to be part of the retractor muscles of the dorsal abdomen^24,31^ and the LSRs organs are repeated in each hemisegment. This simple anatomy makes them well-suited to detect changes in longitudinal muscle stretching and for a coordinated moment-to-moment assessment over multiple segments, similar to other rhythmic behaviours such as larval locomotion^15,39,44^. In tremulation, we might expect the dbd neurons to be stretched when the muscles are relaxed, thus an initial upwards tremulating movement could lead to muscle contraction, followed by a relaxed downwards movement, and so on. The motor neurons and associated muscles responsible for initiating abdominal tremulations are still unidentified; they could be localised within the segment of each LSR or in neighbouring tissues and may rely on feedback from dbd neurons to coordinate their rapid series of contraction and relaxation. Consequently, males deprived of sensory input can only produce reduced and abbreviated tremulations.

In some of cases of patterned movement central pattern generating (CPG) networks have been identified, responsible for generating a basic pattern of CPG-controlled repeated movements with fine-tuning adjustments by the PNS^45,46^. Some central neurons involved in tremulations may potentially share homology with larval neurons participating in locomotor pattern, although there are significant differences in neuromuscular anatomy between larvae and adults. Besides, it is not known if tremulation is also controlled by a CPG. Yet, it is plausible to assume that sensory feedback would be necessary to a CPG to help it retain its efficacy over tremulation bouts lasting 3s or more.

### Proprioception during tremulation

LSR organs have been observed in all insect species studied so far, indicating that our findings are likely relevant for many substrate-borne signalling insects^3,69^. The role of stretch-sensing in body tremulation may extend beyond insects^23,32^. While other body wall neurons could play a role in regulating different aspects of abdominal tremulations, such as their IPI characteristics, our research indicates that nearby ddaC neurons – though they develop complex dendritic arbors on the dorsal body wall of the abdomen but are not associated with muscles^18^ - do not appear to contribute to tremulations.

To produce the characteristic up-and-down oscillatory movement during tremulation, it is expected that both dorsal and ventral muscles are necessary. Future investigations may reveal antagonist ventral muscles that counteract dorsal muscle action and potentially work in tandem with associated sensory neurons to orchestrate the tremulation movements. Adult body wall sensory neurons and ventral abdominal muscles remain understudied areas deserving further exploration^17,47,48^.

### Metamorphosis and neuronal plasticity of stretch-sensing

We investigated the MS ion channels that might be required for dbd neuron stretch function in the tremulations. We found that adult dbd neurons express both *piezo* and *trpa1,* similar to their fly larval counterparts^39^. In larvae, *piezo (*but not *trpa1)* was shown to play a major role in larval dbd stretch function^39^. In adults, by contrast, the depletion of *piezo* in dbd neurons showed no tremulation phenotype. However, the depletion of *trpa1* resulted in significant defects in male abdominal movements, characterised by higher levels of arhythmic movements resembling those observed when dbd neuron function was impaired. Adult dbd neurons therefore appear to require TRPA1 to function in adult stretch-sensing. In larvae each dbd neuron spans a segment antero-posteriorly and it is not inserted in a muscle fiber, but instead associates with epidermal cells at the segmental borders^22^. The remodelling of dbd neurons during metamorphosis is associated with stage-dependent changes in anatomy, adaptation to varied stretch-sensing functions, and a redistribution of roles for MS ion channels. This serves as a striking example of developmental neuronal plasticity. Further findings and the comparison with data on larval dbd will help determine the specific mechanisms of stretch-sensing required for tremulations^11,22,34,39,49^.

### The duration of bouts of tremulation is important for the female’s immobility response

It is unclear which parameters of *Drosophila* tremulation-generated signals are assessed by the female, and which ones convey information that may lead to her immobility response^7,27^. Several characteristics of vibratory signals may be important^3^. Frequency components such as the IPI and/or the amplitude of a signal mediate song recognition in birds, cricket frogs and some Hemiptera^3,50–53^. In *Drosophila*, substrate-borne signals display species-specific and environmentally dependent IPIs, along with amplitude modulation^9,27^. However, results presented here suggest that *Drosophila* tremulation might relate to the signals used by anurans and stoneflies in which the number or the length of the bouts were shown to affect reproductive success^51,54^. Indeed, we found that LSR defects in males impacted the female’s response, likely through the deregulation of bout occurrence and durations. We found that short bouts were less likely to coincide with female immobility, even in wild type courtships. A dependence of the female’s response to the duration of tremulation may relate to the time females take to process that signal and initiate the stop-walking command.

## METHODS

### Resources table

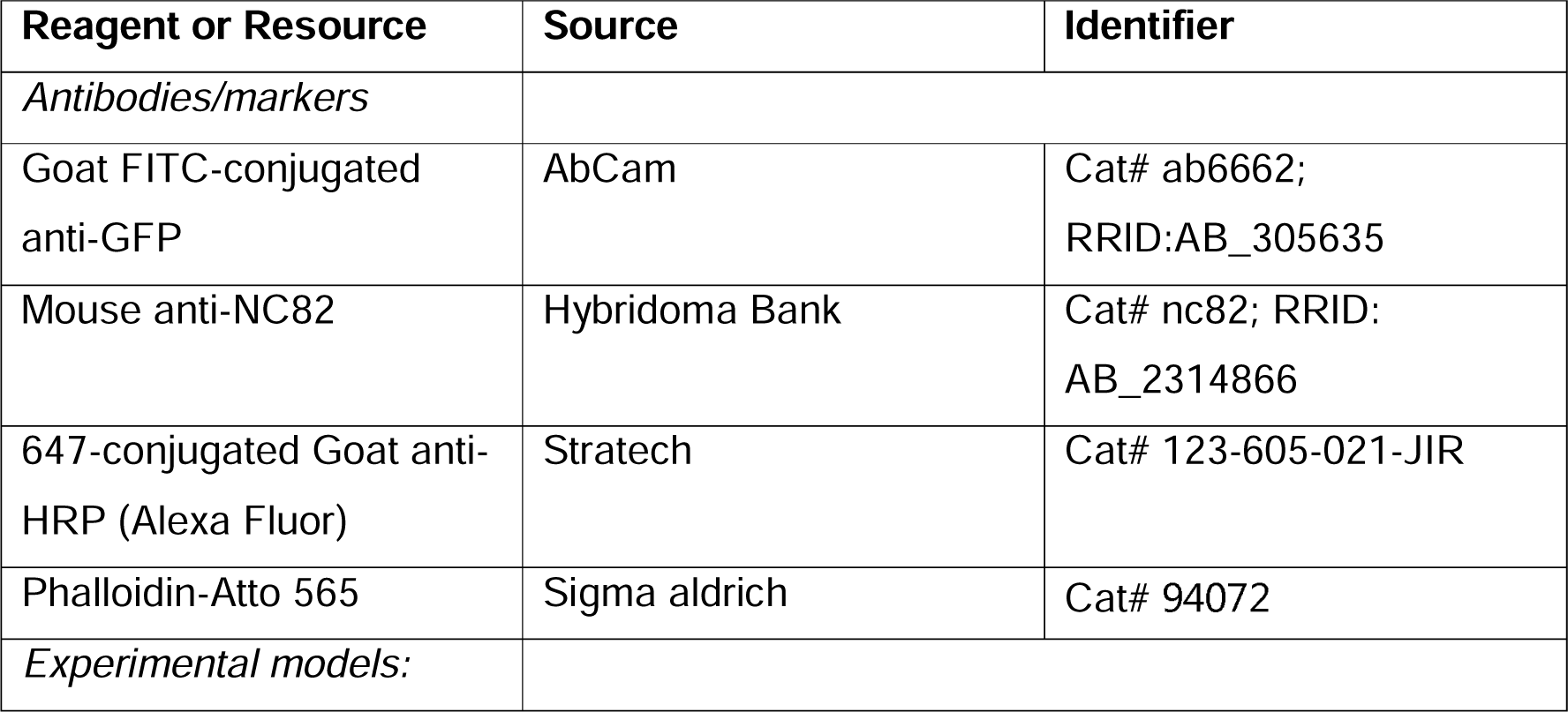
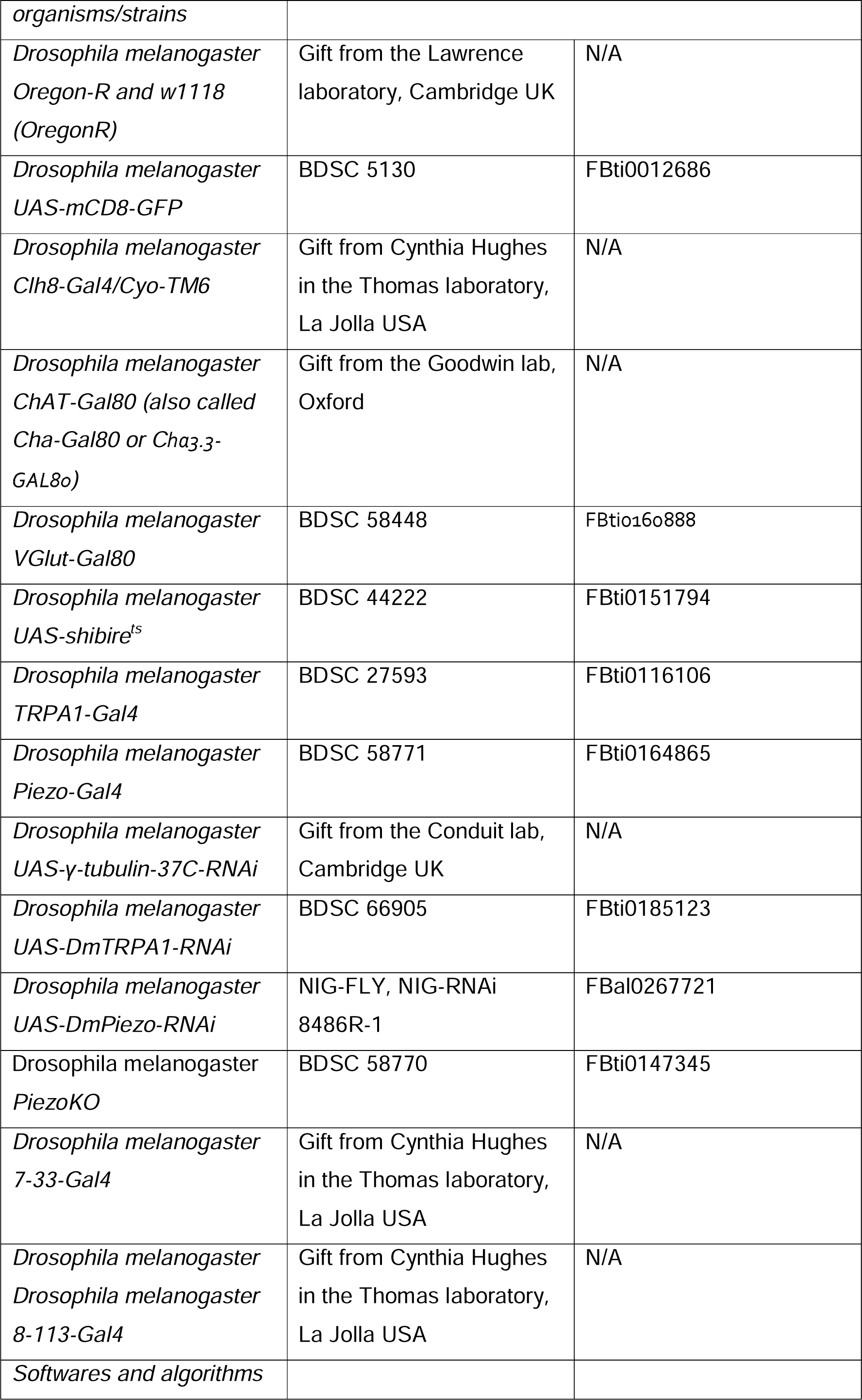
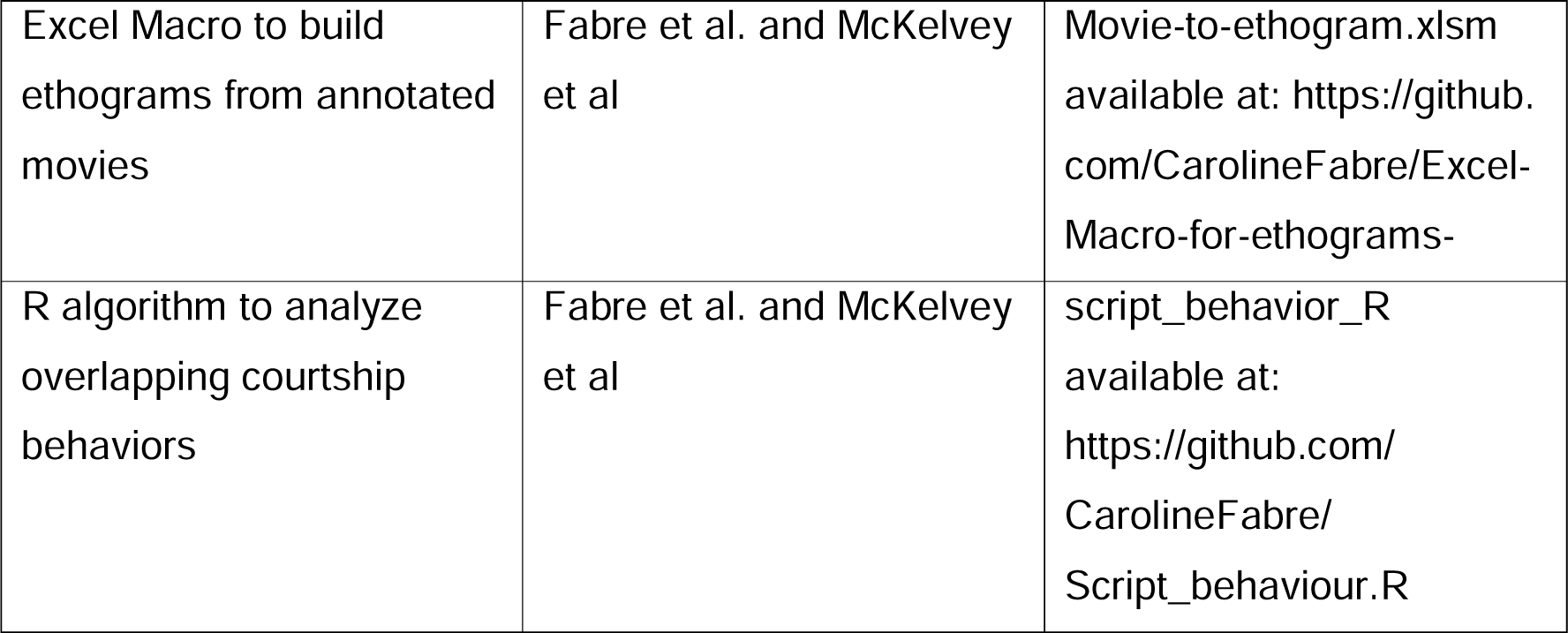

## Materials and Methods

### Fly stocks

Flies were reared under 12:12 hr light:dark cycle, at 23°C and with around 65% humidity. Virgin female/male progeny were collected upon eclosion from the pupal case and tested in courtship assays at ages between 2 and 7 days-old. *dpiezo* null mutant males were obtained by generating females carrying the *dpiezo* knockout (KO) allele and the deficiency Df(2L)Exel7034 (where the entire *dpiezo* genomic region is deleted) on each of the homologous chromosomes 2^20^. Gal4 Stocks were previously backcrossed into a *w1118* (OregonR) background for 3 generations. UAS-shi^ts^ is a dominant-negative variant of dynamin, which can be used to transiently block chemical synaptic transmission at temperature above 29 degrees to silence neurons target by a Gal4 line^25^.

### Immunohistochemistry and microscopy

Adult abdomen dissection and stainings were carried out as described previously^55^. As primary antibodies we used FITC-conjugated anti-GFP (goat serum, Abcam) at 1:500 and 647-conjugated anti-HRP (goat serem, Stratech) at 1:500. Phalloidin (Sigma) was used at 1:1000. Samples were incubated on a slow shaker with the antibodies and the phalloidin for 2 hours at room temperature, or overnight in a cold room. After washes, samples were kept overnight in Fluoromount-G medium (Sigma). Abdomens were mounted flat (with the side of the abdominal bristles upwards or downwards as indicated) on a slide with Fluoromount-G and kept for 2 days at room temperature in the dark for the Fluoromount to solidify. Images were captured on a confocal (Leica SP5) run by LAS AF software. The softwares Fiji and Affinity Designer (Serif) were used to process the .lif and .tif files.

### Behavioral courtship assays and annotations

Behavioural assays were carried out as described previously^9^. Briefly, video-imaging of courting pairs were performed at a temperature of around 23°C. Their behavior was recorded with a 100mm macro lens and a Stingray camera (Allied Vision Technologies; Stadtroda, Germany) and acquired with the Debut Video Capture (Pro Edition) software into a iMac computer. Recording was started at the initiation of courtship and for approximately 600 s, or until copulation occurred. Each pair was tested only once. Behavioural annotations were carried out as described previously^9^. Movies were annotated with semi-automated Annotation software (Peter Brodsky, version 1.3).

### Laser vibrometry

Laser vibrometry of courting pairs were performed at a temperature of around 23°C. Laser vibrometer recordings were conducted on a vibration-damped table in a soundproof room. Flies were filmed and courted when 4 days old upon eclosion from the pupal case. To record, the beam of a laser Doppler vibrometer (Polytec OFV 5000 controller, OFV 534 sensor head; Waldbronn, Germany) was directed perpendicular to the surface of a square of reflective tape (3M, 0.5mm2; Scotchlite, Neuss, Germany) placed in the center onto the surface of the fruit or artificial substrates. Signals were digitised with 12bit amplitude resolution with a PCI MIO-16-E4 card (Analog Devices; Norwood, MA) and with LabView (National Instruments; Austin, TX) on a PC. Signals were transformed into .wav data with Cool Edit Pro (Adobe Systems) or Neurolab software. Oscillograms were analyzed with Amadeus Pro (HairerSoft) and Raven Pro (The Cornell Lab of Ornithology, Bioacoustics Research Program) software.

### Splinkerette PCR (Sp PCR)

All steps followed the spPCR protocol outlined in^36^. Genomic DNA was extracted from 25mg of crushed, adult male *clh8-Gal4* flies using the QIAGEN DNeasy® Blood & Tissue kit (Qiagen). DNA was eluted in 200μl of AE buffer (Qiagen) and purity was confirmed using a NanoDrop 1000 Spectrophotometer (ThermoFisher Scientific). For genomic digestion, a solution was prepared with: 25μl of genomic DNA, 3.5μl of NEB buffer (New England Biolabs), 3.5μl of BSA, 2μl of ddH_2_O and 1μl of BstYI (New England Biolabs). This was incubated for 2hrs at 60°C then heat inactivated for 20min at 80°C. For ligation of digested genomic DNA to splinkerette oligonucleotides, a solution was prepared with: 35μl of digested genomic DNA, 6μl of annealed splinkerette oligonucleotides, 5μl of 10X Ligation Buffer (Roche Diagnostics), 2.5μl of ddH_2_O and 1.5μl of T4 DNA Ligase (Roche Diagnostics). This was incubated for 2hr at 21-23°C. All PCR reactions were conducted on a TProfessional TRIO Thermocycler® (Analytik Jena AG). For spPCR Round 1, a reaction mix was prepared with: 10μl of ligated genomic DNA, 8.25μl of ddH_2_O, 12.5μl of 2X ThermoFisher Scientific® Phusion® High-Fidelity PCR Master Mix with HF Buffer (New England Biolabs) and 0.5μl each of primers 10μM SPLINK#1 (Sigma) and 10μM 5’SPLINK#1-GAWB (Sigma). For spPCR Round 2, a reaction mix was prepared with: 1μl of Round 1 PCR, 12.5μl of the PCR Master Mix used in Round 1, 11.5μl of ddH_2_O and 0.5μl each of primers 10μM SPLINK#2 (Sigma) and 10μM 5’SPLINK#2-GAWB (Sigma). The Round 2 PCR product was run on a 1% agarose gel, made using 1g of UltraPure^TM^ Agarose (ThermoFisher Scientific) in 100ml of 1X TBE buffer (ThermoFisher Scientific) with 10μl of Sybr safe DNA stain (ThermoFisher Scientific). 5μl of Round 2 PCR product was loaded with 1μl of DNA loading buffer (Bioline). 3μl of Hyperladder 1kb (Bioline) was loaded for size comparison. Electrophoresis was performed at 80V for 30min in 1X TBE buffer. Gels were imaged using a Safe Imager^TM^ 2.0 Blue Light Transilluminator (ThermoFisher Scientific). The PCR band was gel-extracted using the QIAquick® Gel Extraction Kit (Qiagen) and sequencing was performed by Source BioScience (Cambridge, UK).

## Acknowledgements

We thank Paul Conduit, Peter Lawrence and Jose Casal for discussions and feedback on the data and manuscript. This project has received funding from the European Union’s Horizon 2020 research and innovation programme under the Marie Skłodowska-Curie grant agreement No 945298-ParisRegionFP. C.C.G.F is a Fellow of Paris Region Fellowship Programme supported by the Paris Region. This work was also supported by the Wellcome Trust (P.L.’s grant number 107060/Z/ 15/Z) and a Sir Isaac Newton Trust Research Grant (16.24(i) (C.C.G.F). The work benefited from use of the Imaging Facility, Department of Zoology, supported by a Sir Isaac Newton Trust Research Grant (18.07ii(c)).

## Figure legends

**Figure S1.**
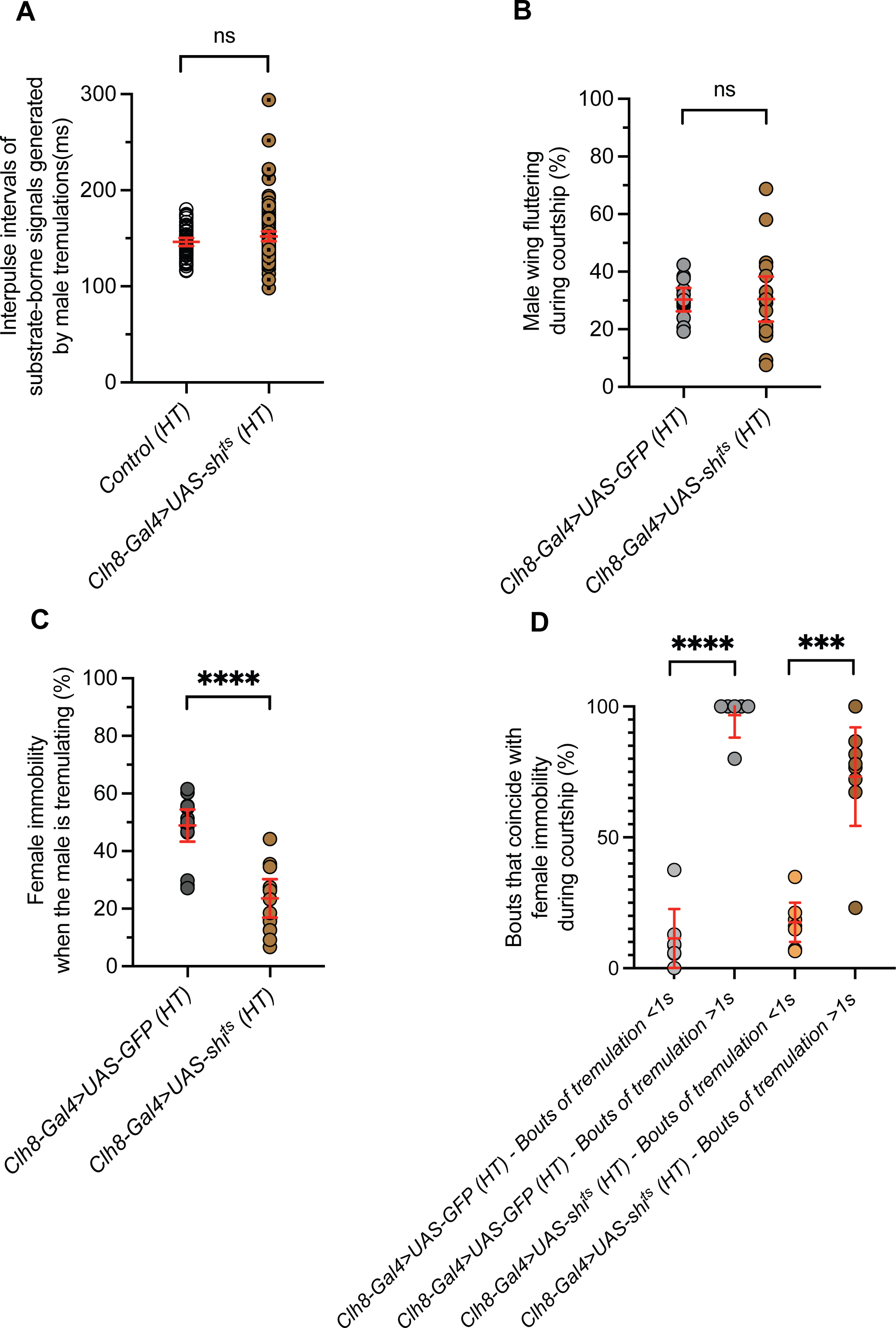
Comparison of male and female behaviours during courtship of pairs including a *Clh8-Gal4>UAS-shi^ts^* or a control *Clh8-Gal4>UAS-GFP* male at restrictive temperature. Same pairs as in Figure 1. In all pairs females are wild-type OregonR. Pairs were monitored at high temperature (HT). **(A)** Values of interpulse intervals (IPIs, in ms) that characterise substrate-borne vibratory signals are similar, whether they are generated by tremulations of control or *Clh8-Gal4>UAS-shi^ts^*males at high temperature. **(B)** shows another courtship behaviour, wing fluttering, is displayed at a similar percentage of the courtship time by both control and *Clh8-Gal4>UAS-shi^ts^*males at high temperature. **(C)** The percentage of time females remain immobile during male tremulations is significantly decreased when they are paired with *Clh8-Gal4>UAS-shi^ts^* males at high temperature. **(D)** The percentage of tremulation bouts that coincide with female immobility is assessed depending on the duration of the bouts, both in pairs including *Clh8-Gal4>UAS-GFP* males and in pairs including *8-Gal4>UAS-shi^ts^* males. In both types of pairs, bouts with a duration higher than 1s (>1s) coincide mainly with female immobility, while bouts with a duration lower than 1s (<1s) coincide significantly less with female immobility.

**Figure S2.**
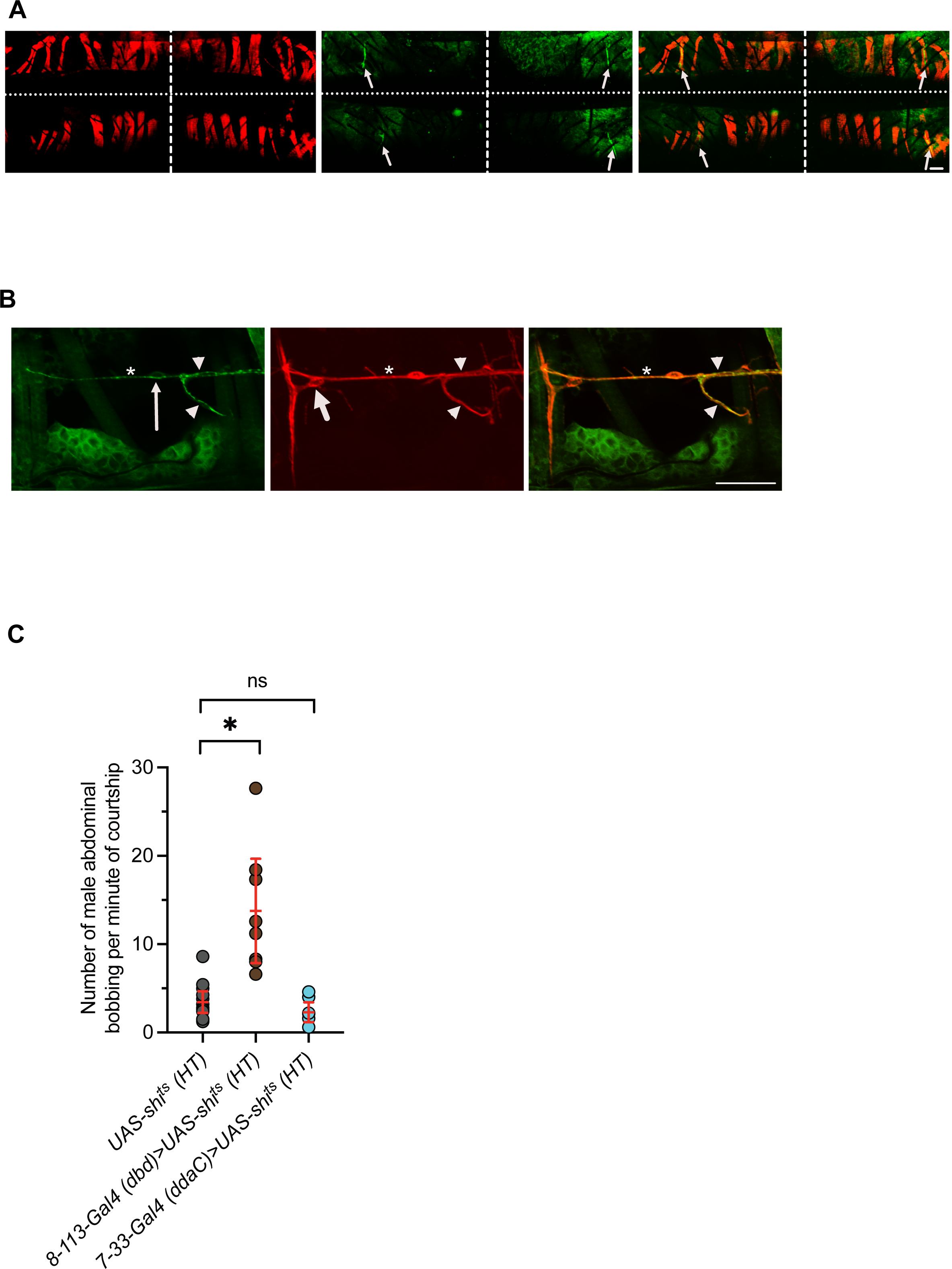
Expression patterns and bobbing behaviours characterising other selected Gal4 lines. **(A)** Staining of segments 2 and 3 of the abdomen of a male carrying *8-113-Gal4>UAS-mGFP* shows *8-113-Gal4* drives expression in the dbd neurons (anti-GFP, green; arrows) in each hemisegment (horizontal dashed line separates the segments 2 and 3; vertical dashed line indicates dorsal midline). Muscles are marked using phalloidin (red). Note, some muscle fibers were separated and dispersed during the dissection. **(B)** Staining of the third left hemisegment of the abdomen of a male carrying *7-33-Gal4>UAS-mGFP* shows expression in the (larval class IV) ddaC neuron cell body (thin arrow; anti-GFP, green) and neurites (green). ddaC is recognisable by its localisation immediately proximate to the dbd neuron (large arrow; anti-HRP, red), and by its cell body localised onto the nerve (anti-HRP, red) on which its axon bundles (asterix). Part of the ddaC neuron dendritic arborization (arrowheads) can be seen towards the dorsal midline that is localised to the right. Anterior up, Posterior down.

**Figure S3.**
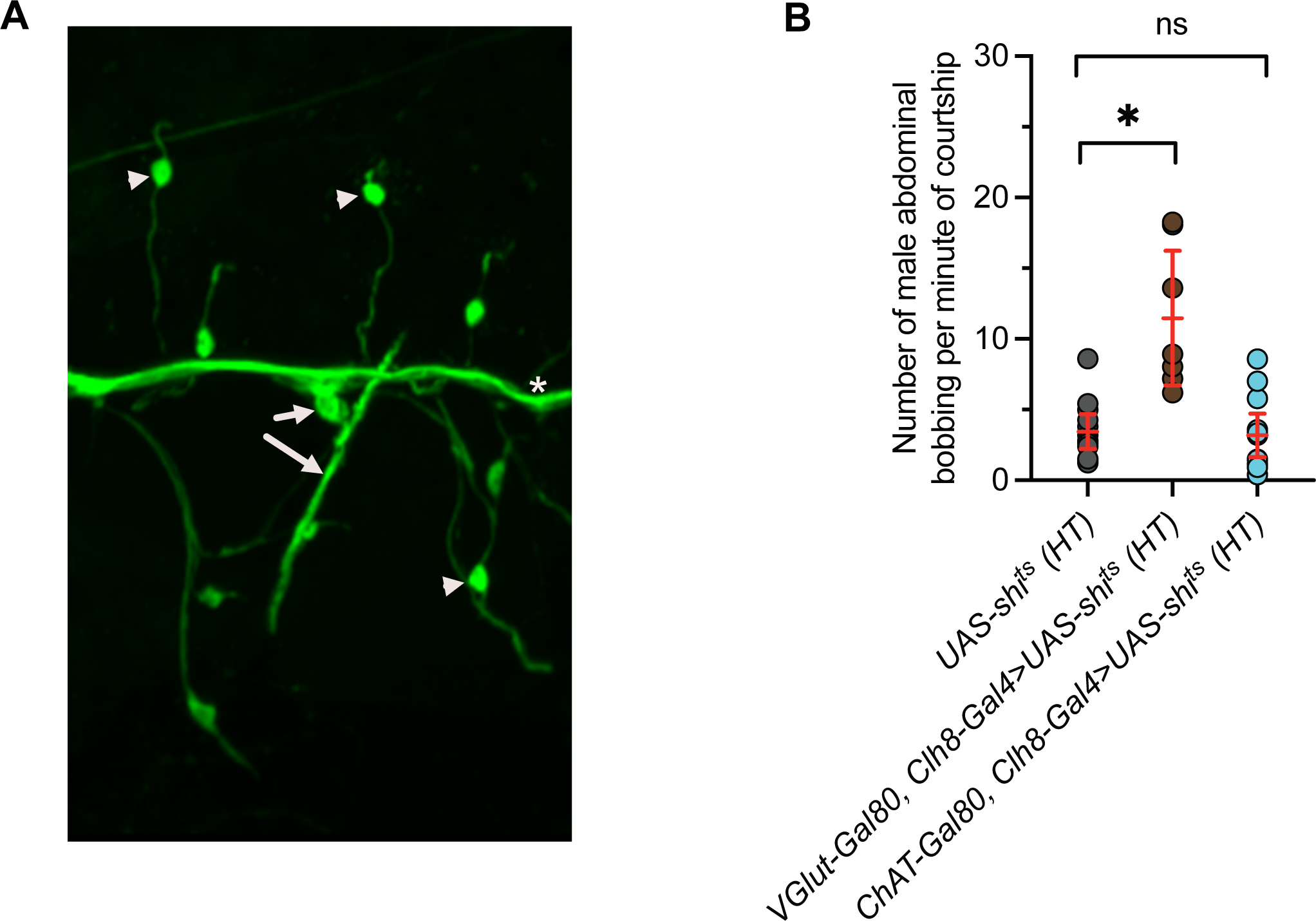
Effect of restricting the expression of *Clh8-Gal4>UAS-shi^ts^* in males during courtship. (**A**) ChAT-Gal4>UASmGFP shows expression in adult dbd neurons (arrows, anti-GFP, green), indicating that dbd neurons express choline acetyltransferase (ChAT) and are cholinergic. Other cholinergic neurons visible are the mechanosensory neurons of abdominal bristles (arrowheads). (B) In all cases females are OrR. All courtship were filmed at HT. Controls are males carrying the *UAS-shi^ts^* contructs only. Using a promoter to express Gal80 specifically in glutamatergic neurons (*VGlut-Gal80*) had no effect (2^nd^ column, brown). However, expressing the Gal4 inhibitor Gal80 specifically in cholinergic neurons (*ChaT-Gal80*) significantly reduced bobbing in *Clh8-Gal4>UAS-shi^ts^* males (3^rd^ column, light blue).

**Figure S4.**
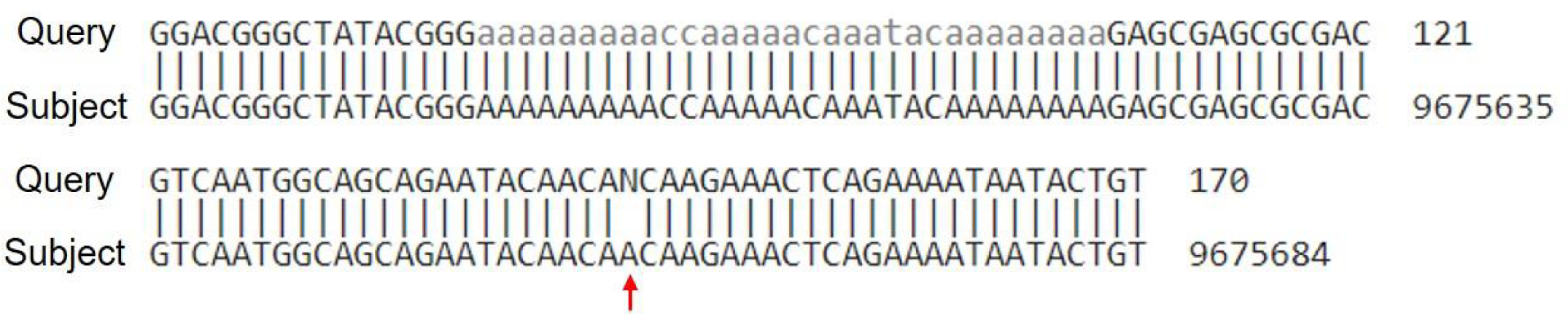
Splinkerette-PCR identified *furry* as a candidate gene for *P{GawB}Clh8* insertion location. There is high local sequence similarity between the sequence flanking *P{GawB}Clh8* and the *furry* gene on chromosome 3L. Result was generated by NCBI BLAST. Alignment block showing our query sequence along the top and subject sequence underneath, which is a portion of the *furry* gene. Numbers on the right indicate nucleotide (nt) position. Identical base match is represented by a line. In this local region, 108/109 identities match (99%), indicating high local sequence similarity. The non-matching pair is indicated by a red arrow.

**Figure S5.**
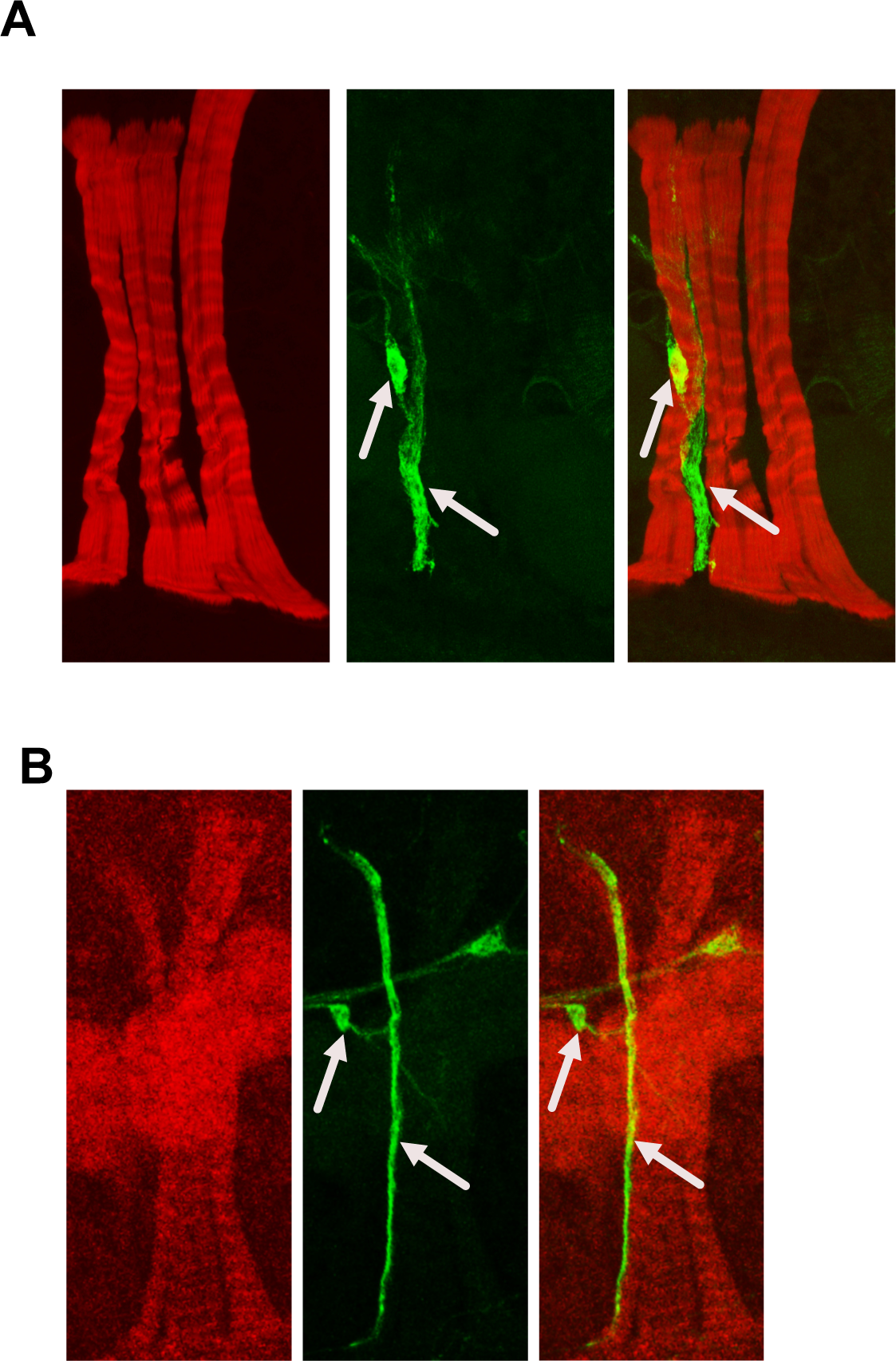
Expression of the Gal4 lines *dPiezo-Gal4* and *dTrpa1-Gal4* in the dbd neurons. Gal4 lines were combined with *UAS-mGFP* and show, in a third hemisegment of an abdomen, that **(A)** *dTrpa1-Gal4* and **(B)** *dPiezo-Gal4* both drive expression in the dbd neuron cell body and neurites (arrows, anti-GRP, green). Muscles are labelled using Phalloidin (red).

## Notes

### Competing Interest Statement

The authors have declared no competing interest.

### Summary of Updates

I amended the order of the authors in the BioRxiv submission form as I had entered it in the wrong order at submission, I apologise for this. Everything else is similar to the original submission. Many thanks. Caroline Fabre

